# A checkpoint roadmap for the complex cell division of Apicomplexa parasites

**DOI:** 10.1101/104646

**Authors:** Carmelo A. Alvarez, Elena S. Suvorova

## Abstract

The unusual cell cycles of Apicomplexa parasites are remarkably flexible with the ability to complete cytokinesis and karyokinesis coordinately or postpone cytokinesis for several rounds of chromosome replication are well recognized. Despite this surprising biology, the molecular machinery required to achieve this flexibility is largely unknown. In this study, we provide comprehensive experimental evidence that apicomplexan parasites utilize multiple Cdk-related kinases (Crks) to coordinate cell division. We determined that *Toxoplasma gondii* encodes seven atypical P-, H-, Y- and L- type cyclins and ten Crks to regulate cellular processes. We generated and analyzed conditional tet-OFF mutants for seven TgCrks and four TgCyclins that are expressed in the tachyzoite stage. These experiments demonstrated that TgCrk1, TgCrk2, TgCrk4 and TgCrk6, were required or essential for tachyzoite growth revealing a remarkable number of Crk factors that are necessary for parasite replication. G1 phase arrest resulted from the loss of cytoplasmic TgCrk2 that interacted with a P-type cyclin demonstrating that an atypical mechanism controls half the *T. gondii* cell cycle. We showed that *T. gondii* employs at least three TgCrks to complete mitosis. Novel kinases, TgCrk6 and TgCrk4 were required for spindle function and centrosome duplication, respectively, while TgCrk1 and its partner TgCycL were essential for daughter bud assembly. Intriguingly, mitotic kinases TgCrk4 and TgCrk6 did not interact with any cyclin tested and were instead dynamically expressed during mitosis indicating they may not require a cyclin timing mechanism. Altogether, our findings demonstrate that apicomplexan parasites utilize distinctive and complex mechanisms to coordinate their novel replicative cycles.

## INTRODUCTION

Obligate intracellular parasites of Apicomplexa phylum are responsible for many important diseases in humans and animals, including malaria, toxoplasmosis and cryptosporidiosis. Severity of the disease is tightly linked to parasite burden, and currently, the most successful therapies block parasite proliferation. Apicomplexan parasites use flexible mechanisms to replicate that are different than those of their hosts. In endodyogeny each round of duplication is completed with assembly of two internal daughters [1]. Alternatively, multiple buds can be formed inside of the mother or emerge from its surface in the processes called, endopolygeny and schizogony, respectively [2]. *Toxoplasma gondii* undergoes endodyogeny in intermediate host stages, but replicates by endopolygeny in the definitive feline host. Fundamental differences between division modes are embedded in the features of the apicomplexan cell cycle comprised of two chromosome cycles [2, 3]. During nuclear cycles, chromosomes are replicated and segregated without budding, while each round of chromosome replication in the budding cycle leads to production of the daughter parasites. Ultimately, the number of nuclear cycles determines the scale of the parasite progeny. We recently showed that the ability to switch between chromosome cycles is partially linked to the unique bipartite structure of the *T. gondii* centrosome [3]. Weakening or separation of the outer centrosomal core that controls budding favors the nuclear cycle, while the strong association of the outer core with the inner core promotes cytokinesis and the budding cycle [3].

Endodyogeny of *T. gondii* tachyzoites represents one the simplest modes of replication and important cell cycle transition points where potential checkpoints may operate have been defined [2, 4–19]. The G1 phase of *T. gondii* endodyogeny comprises half of the division cycle, and like other eukaryotes, canonical housekeeping tasks preparing for S phase commitment are performed in the apicomplexan G1 period [4, 9, 13, 15, 18, 19]. Evidence also indicates that cell cycle exit to form dormant developmental stages as well as drug-induced dormancy is controlled by mechanisms acting at the transition from M/C into early G1 [4, 20, 21]. Duplication of the centrosome marks the transition from G1 to S phase [3, 6, 15, 17], and we have defined some of the components of this critical transition in *T.gondii* that should be targets of a G1/S checkpoint mechanism [17]. A peculiar feature of apicomplexan replication is the short (or absent) G2 phase [1, 2] that is thought to be marked by natural S phase populations that possess partially duplicated genomes (1.7-1.8N DNA) [1, 13]. Resolving the molecular basis for this important transition should help solve the mystery of these unusual DNA distributions. During the apicomplexan mitosis numerous specialized structures are replicated, built or converted in precise order to produce healthy infectious daughters (for review see [2, 22]). Our previous work and the studies of others have established that duplication of the bipartite centrosome is coordinated with division of the centrocone that holds the mitotic spindle [3, 6, 10, 23], and also with the replication and segregation of the bundled centromeres [23, 24], which in turn, is synchronized with assembly of the basal and apical complexes of the future daughter [23–28]. The temporal-spatial coordination of these overlapping processes likely requires similarly complex regulatory machinery, for which the molecular basis is still largely unknown.

In eukaryotes, cell cycle progression is governed by the activity of cyclin-dependent kinases (Cdks) and their regulatory cofactors, cyclins [29, 30]; dynamic expression of the latter provides clockwise control of Cdk function. Cdk4/6-cyclin D complexes support the progression of G1 phase, and Cdk2 complexes with cyclins E, A and B govern the progression and fidelity of DNA replication in S phase and chromosome segregation in mitosis [31]. Cdks functions were originally thought to be restricted to cell cycle regulation, however, today we understand that activated Cdk-cyclin complexes are master regulators of such major biological processes as transcription, RNA processing, translation and development [30]. Extrapolating current models of cell cycle checkpoints that involve Cdk-cyclins to eukaryotes in general is challenging, as there are many branches of the eukaryotic tree where cell division is quite unusual and the molecular controls are likely to be different [29]. This includes the large group of obligate intracellular parasites from the phylum Apicomplexa. Mining the initial genomes of important disease causing apicomplexans has revealed major differences [1, 32–34] characterized by the reduction of components and also the complete absence of the key regulatory elements, including canonical cyclins [35], major cell cycle Cdks [33, 34] and their immediate downstream effectors [1]. The lack of conserved cell cycle factors of higher eukaryotes indicates there are significant changes in the cell cycle molecular machinery of these ancient protozoa. Here we describe the first comprehensive study of Cdk-related kinases (Crks) and cyclins in *T. gondii*. Using genetic approaches, we have analyzed the function of seven TgCrks and four TgCyclins and uncovered major cell cycle TgCrks controlling the replication of the tachyzoite stage of *T. gondii*. Our results demonstrate that, unlike the traditional eukaryotic cell cycle, the intricate division of apicomplexan parasites is regulated by multiple essential Crks acting independently at several critical transitions and in unusual spatial contexts.

## RESULTS

### *Toxoplasma gondii* possesses multiple divergent Cdk-related kinases

To determine the core Cdk-cyclin complexes that regulate division in *T. gondii*, we systematically searched the parasite genome and identified ten genes encoding a kinase domain that included a cyclin-binding sequence (C-helix) (S1 Fig) [30]. Following standard convention, we named these factors Cdk-related kinases (Crks) until cyclin-dependent activation of the kinase can be established [36, 37]. *T.gondii* Crks (TgCrks) were a diverse group of proteins ranging from 34 to 212kDa (S1A Fig) that also varied in mRNA abundance (S1B Fig). *T. gondii* transcriptome data (ToxoDB) indicated that eight TgCrks were expressed in tachyzoites and/or bradyzoites, and mRNA profiles for two kinases, TgCrk2-L1 and TgCrk5-L1 indicated they were restricted to the definitive or environmental life cycle stages; merozoite and/or sporozoite (ToxoDB). The profiles of TgCrk4, TgCrk5 and TgCrk6 mRNAs were cyclical in tachyzoites (S1B Fig) [4], which was confirmed at the protein level for TgCrk4^HA^ and TgCrk6^HA^ by endogenous epitope tagging (S2 Fig). The dynamic cell cycle regulation of the three TgCrk factors differs significantly from the typical constitutive expression of Cdks of studied model eukaryotes [38, 39].

Phylogenetic analysis of TgCrks defined the general Cdk families present in the Apicomplexa phylum [34]. Putative Crks from *T. gondii* and *P. falciparum* [33, 34] were compared to the ancestral free-living unicellular eukaryote *Chromera velia*, and the extensively studied Cdks of human cells. The analysis sorted ten TgCrks into six general phylogenetic clades with three clades restricted to the superphylum Alveolata (Fig 1, pink shade): TgCrk3, TgCrk4, TgCrk5 and TgCrk6 were either apicomplexan or coccidian adaptations and two pairs TgCrk3/TgCrk8, and TgCrk4/TgCrk6 likely had a common ancestor. Kinases in three other clades were shared with a recognizable higher eukaryotic counterpart that included several kinases known to regulate cell cycle and gene expression. Specifically, TgCrk1 grouped with the Cdk11 family kinases that regulate mRNA synthesis and maturation (Fig 1, yellow shade), while TgCrk7 was similar to the Cdk-activating kinase (CAK), HmCdk7 (Fig 1, green shade). Lastly, TgCrk2 clustered with the family of eukaryotic cell cycle regulators HmCdk1/2/3 and neuronal HmCdk5 (Fig 1, blue shade).

**Fig 1.**
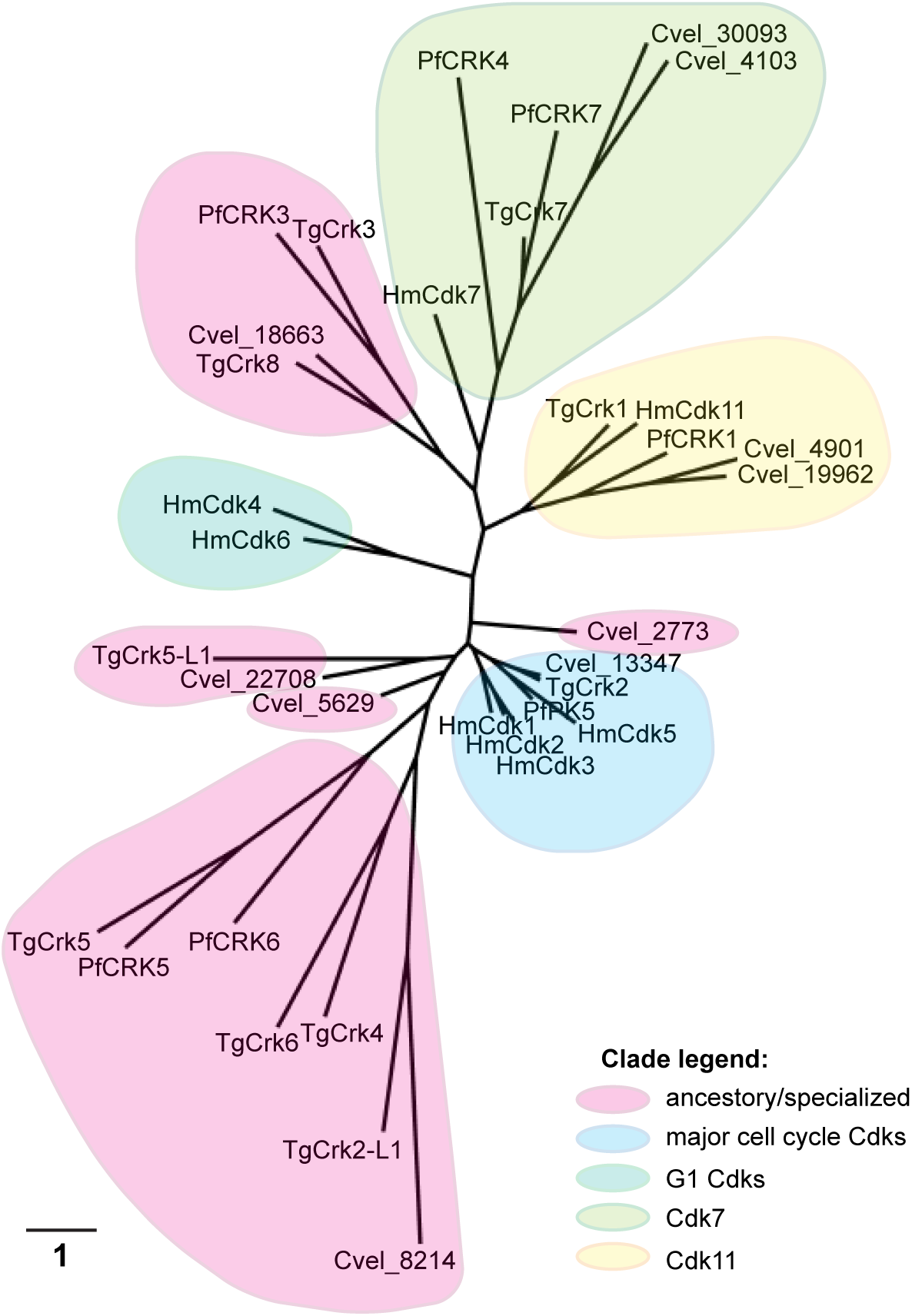
**Phylogenetic analysis of *T. gondii* Crks**. Protein sequences of Cdk-related kinases from *T. gondii, P. falciparum, C. velia*, and Cdks from human cells were analyzed in Phylogeny.fr software. The results show that three Cdk families are shared between Alveolates and higher eukaryotes: *T. gondii* TgCrk2 was grouped with cell cycle Cdk1/2 family kinases (blue shadow), TgCrk1 was clustered with transcriptional kinases of Cdk11/12 family (yellow shadow) and TgCrk7 appeared distantly related to CAK complex component Cdk7 (green shadow). Note that *T. gondii* does not encode canonical G1 Cdk4/6 family (teal shadow). Clades that lack higher eukaryotic counterpart were labeled as ancestral/specialized and indicated with pink shadow. The majority of *T. gondii* Crks were clustered in ancestral clades.

### Conditional knockdown of *Toxoplasma gondii* Cdk-related kinases

To define the function of TgCrks, we constructed tet-OFF conditional knockdown mutants by replacing native promoter with a tetracycline-regulatable promoter in the Tati-RHΔ*ku80 strain* [40, 41]. Each kinase was concurrently tagged with a 3xHA-epitope fused to the N-terminus (S1D Fig, diagram). We successfully generated tet-OFF mutants for seven TgCrks demonstrating that promoter replacement and N-terminal HA-tagging were remarkably tolerated in the tachyzoites (Fig 2). The only other TgCrk factor expressed in tachyzoites is TgCrk5, which is the subject of another project and was not studied here (Naumov and White, personal communication). Immunofluorescence assays (IFA) of ^HA^TgCrks determined that most of these kinases were predominantly nuclear in tachyzoites (Fig 2, -ATc conditions) with the exception of ^HA^TgCrk2, which was expressed throughout the cell, and ^HA^TgCrk4, which was exclusively localized to the cytoplasm. Despite sharing the same tet-OFF promoter, individual ^HA^TgCrks showed a wide range of protein abundance indicating there are major post-transcriptional mechanisms controlling TgCrk levels in tachyzoites (S1E Fig).

**Fig 2.**
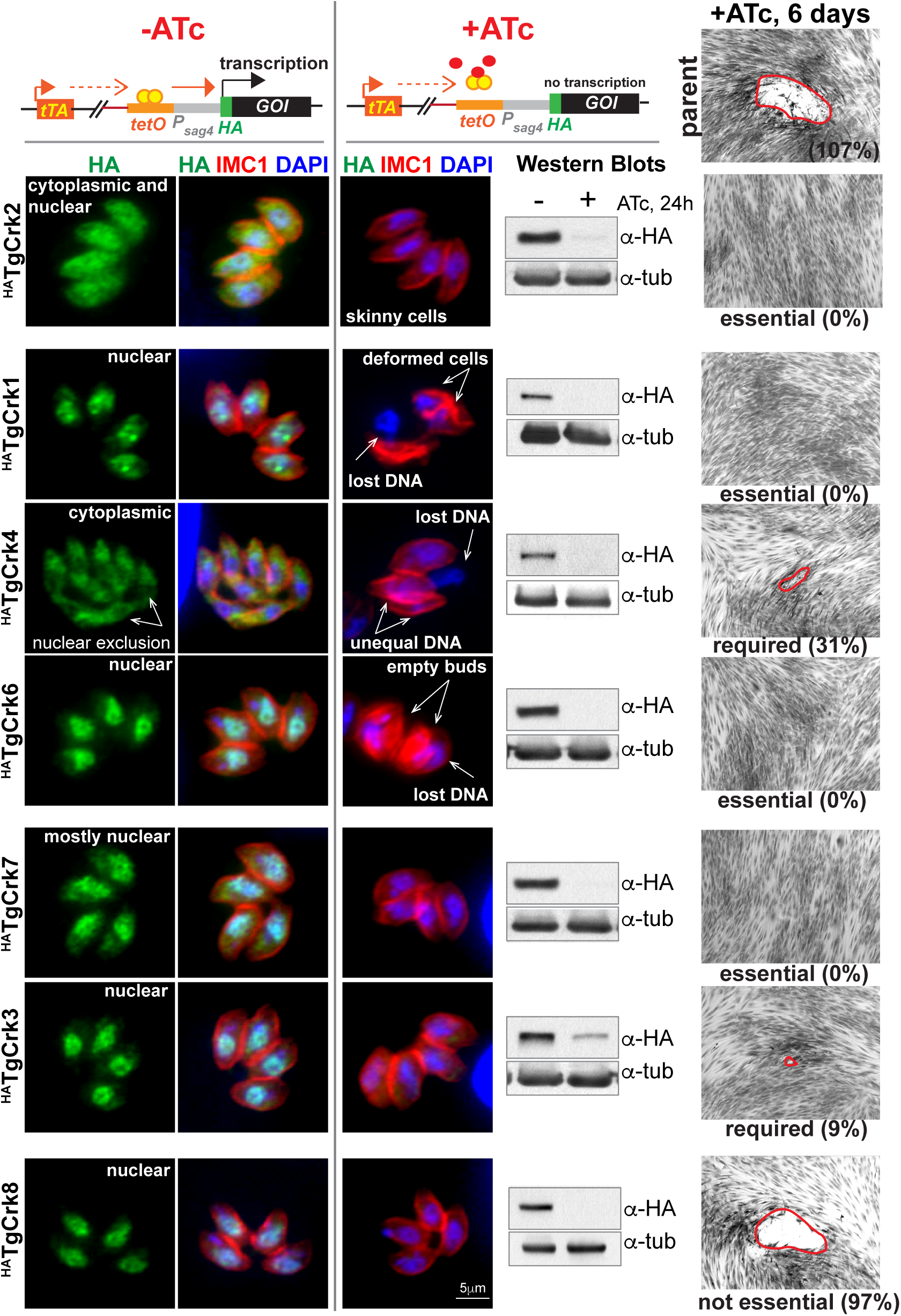
**Conditional expression of *T. gondii* Crks**. The upper schematic illustrates anhydrotetracycline (ATc) mediated control of gene expression in the tet-OFF system. In the absence of ATc the tetracycline transactivator (tTA) binds to the tet-operator (tetO) maintaining the active transcription of the gene of interest (GOI). Downregulation of the GOI product is achieved by transcriptional repression with ATc that prevents tTA binding to tet-operators incorporated into the tet-OFF promoter of the GOI. IFA images on the left show predominant localization of the epitope-tagged kinases (α-HA, green) relative to nuclear staining (DAPI, blue) and IMC compartment (IMC1, red). Downregulation of TgCrk expression after 24 h treatment with 1µg/ml ATc was verified for seven kinases by IFA and Western Blot analysis (α-Tubulin A staining was used as a loading control). The essentiality of each kinase was tested by ability of TgCrk tet-OFF mutant parasites to form plaques after 6 days with 1µg/ml ATc (representative DIC images on the right). Parent Tati-RHΔ*Ku80* strain was included as positive control (top DIC image). Percentage number indicated below each DIC image represents the number of plaques relative to the -ATc condition for each tet-OFF mutant. All analyzed kinases were either essential or required for tachyzoite growth, with the exception of TgCrk8. Downregulation of TgCrk1, TgCrk2, TgCrk4 and TgCrk6 led to major cell cycle defects that are indicated in the IFA images.

The ^HA^TgCrk proteins were all successfully down regulated by a 24 h incubation with 1µg/ml anhydrotetracycline (ATc) (Fig 2, IFA and Western blot analysis). Exploiting the ATc-induced conditional knockdown, we determined by plaque growth assay (Fig 2) that four TgCrks were essential, two were required for tachyzoite growth and only TgCrk8 was dispensable. Quantitative growth rates (24 h) for the TgCrk4 tet-OFF mutant with or without ATc treatment confirmed that this factor was required, but not essential for tachyzoite growth (S1F Fig). IFA analysis determined that ATc-induced growth arrest of TgCrk3 and TgCrk7 tet-OFF mutants was cell cycle independent (Fig 2, IFA +ATc) consistent with the potential role in regulating general metabolic processes. Recent studies of TgCrk7, and its *P. falciparum* ortholog Pfmrk, implicated a role in transcriptional regulation [42, 43] and PfCRK3 kinase complexes were associated with chromatin-dependent regulation of the gene expression [37]. Based on parasite morphology, the loss of TgCrk2 appeared to block tachyzoites in the G1 phase and knockdown of TgCrk1, TgCrk4 or TgCrk6 resulted in extensive mitotic and cytoskeletal defects, consistent with growth arrest in the S/M/C half of the cell cycle. These four kinases were selected for further characterization in this study.

### Determining the function of cyclins in *Toxoplasma gondii*

In general cyclins are poorly conserved, although the presence of a cyclin box and one or more destruction motifs permits these genes to be identified by genome mining. Utilizing this approach we identified seven novel cyclin factors in the *T. gondii* genome (S3A Fig) [35]. Only cyclins related to P-, H-, L- and Y- types were found, while no canonical A-, B-, D-, and E- types, that are vital to higher eukaryotic cell division, were identified [44]. Based on the extensive transcriptome data (see ToxoDB.org), five of the seven TgCyclins appeared to be expressed in tachyzoites (log^−2^ RMA value higher then 6, S3A Fig). To determine expression and localization of TgCyclins in tachyzoites, we epitope tagged the C-terminus of of TgCycH, TgCycL, TgCycY with a 3xHA-epitope by genetic knock-in. IFA analysis showed that TgCycH^HA^, TgCycL^HA^ and TgCycY^HA^, were moderately expressed and localized to the nucleus in tachyzoites (Fig 3A), and remarkably, TgCycY^HA^ was the only oscillating cyclin with peak expression in the G1 phase (Fig 3B). To visualize the lower abundant cyclin, TgPHO80, we engineered transgenic parasites ectopically expressing TgPHO80 that was N-terminally tagged with a 3xmyc epitope fused to a FKBP destabilization domain (DD^myc^), which permits conditional expression using the small molecule Shield 1 [15, 45]. After 3 h stabilization with Shield 1 (100nM), DD^myc^TgPHO80 was found in large cytoplasmic speckles (Fig 3A), which was the only cyclin exclusively localized to the tachyzoite cytoplasm.

**Fig 3.**
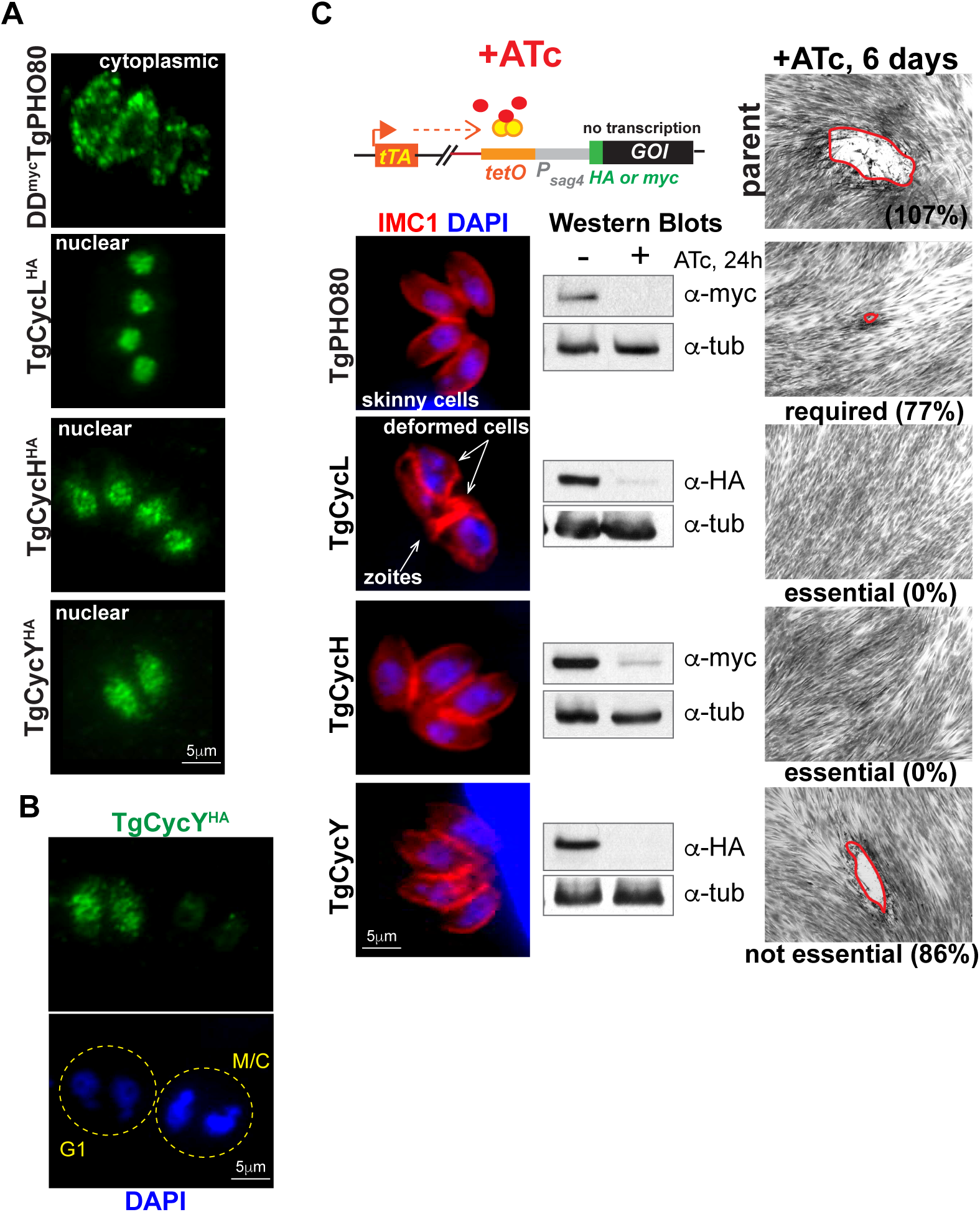
**Conditional expression of *T. gondii* cyclins in the tet-OFF model**. (A) IFA analysis of TgCyclins localization. Tagged at the genomic locus TgCycH^HA^, TgCycL^HA^ and TgCycY^HA^ were visualized using α-HA antibody (green). Due to low native expression level, localization of ectopically expressed DD^myc^TgPHO80 is shown after a 3 h induction with 100nM Shield1. (B) TgCycY^HA^ is dynamically expressed in G1 stage. Yellow dotted line indicates individual vacuoles. The cell cycle stage was determined based on intensity of the nuclear staining (DAPI, blue). (C) Transgenic tet-OFF clones of TgCyclins were established with alternative 3xHA or myc tags as described in Material and Methods and in the S1D Fig. IFA images show a representative vacuole after 24 h growth with 1µg/ml ATc. Parasite shape and nucleus were visualized with α-IMC1 (red) and DAPI (blue) staining, respectively. ATc-induced downregulation of the TgCyclins was confirmed by Western blot analysis using α-Tubulin A staining as a loading control. Results of the plaque assays are shown after 6 days growth with 1µg/ml ATc (DIC panel on the right). The percentage is the number of plaques formed relative to no ATc condition. The phenotype of TgPHO80 and TgCycL deficient parasites suggested factors involvement in the cell cycle regulation.

Utilizing a similar knockdown approach as was used for the TgCrks (S1D Fig), we successfully converted by genetic knock-in four TgCyclin genes to tet-OFF mutant alleles: TgPHO80, TgCycH, TgCycL and TgCycY (Fig 3C, Western Blot analysis). ATc-knockdown of TgPHO80 slowed the rate of replication (S3C Fig) and increased the fraction of parasites with a single centrosome (76.5% ± 9.8 vacuoles after 48 h with 1µg/ml ATc compared to 49% ± 1 vacuoles in -ATc conditions) indicating the G1 phase was lengthened by the loss of TgPHO80. The ATc-induced depletion of nuclear cyclins, TgCycL and TgCycH, caused tachyzoite growth arrest. TgCycL deficiency led to mitotic death and knockdown of TgCycH resulted in a quick non-specific growth arrest (Fig 3C). The conditional knockdown of TgCycY with ATc demonstrated this cyclin was not essential for tachyzoite growth (Fig 3C and growth quantification in S3C Fig).

### TgCrk2 and P-type cyclin regulate the tachyzoite G1 phase

The conditional knockdown of the tet-OFF TgCrk2 mutant appeared to cause a cell cycle arrest in G1, which we confirmed by analyzing centrosome duplication. As expected, the majority (~80%) of TgCrk2-depleted parasites (+ATc) possessed a single centrosome (Fig 4A) indicative of G1 phase arrest. Since canonical G1 cyclins (e.g. D- and E- type) [29] were absent from the *T. gondii* genome, it was of particular interest to identify the *T. gongii* cyclin that binds TgCrk2. We genetically engineered a series of dual epitope-tagged strains by first knocking-in 3xHA epitope tag into the *TgCrk2* gene followed by stable ectopic expression of four different TgCyclins that were epitope tagged with 3xmyc (see S1 Table for list of transgenic strains and Material and Methods for production details). TgCrk2 protein complexes were affinity purified with antibody against epitope tag and then probed with alternative antibody to define the interacting cyclin. We detected a weak interaction between TgCrk2 and TgCycH (Fig 4B) confirming previous yeast two-hybrid screens [46], however, TgCrk2^HA^ formed the most abundant complexes with cyclin TgPHO80-1 (Fig 4B), which corroborates the G1 phenotype we observed for the TgPHO80 tet-OFF mutant following ATc-knockdown (Fig 3C and S3C Fig).

**Fig 4.**
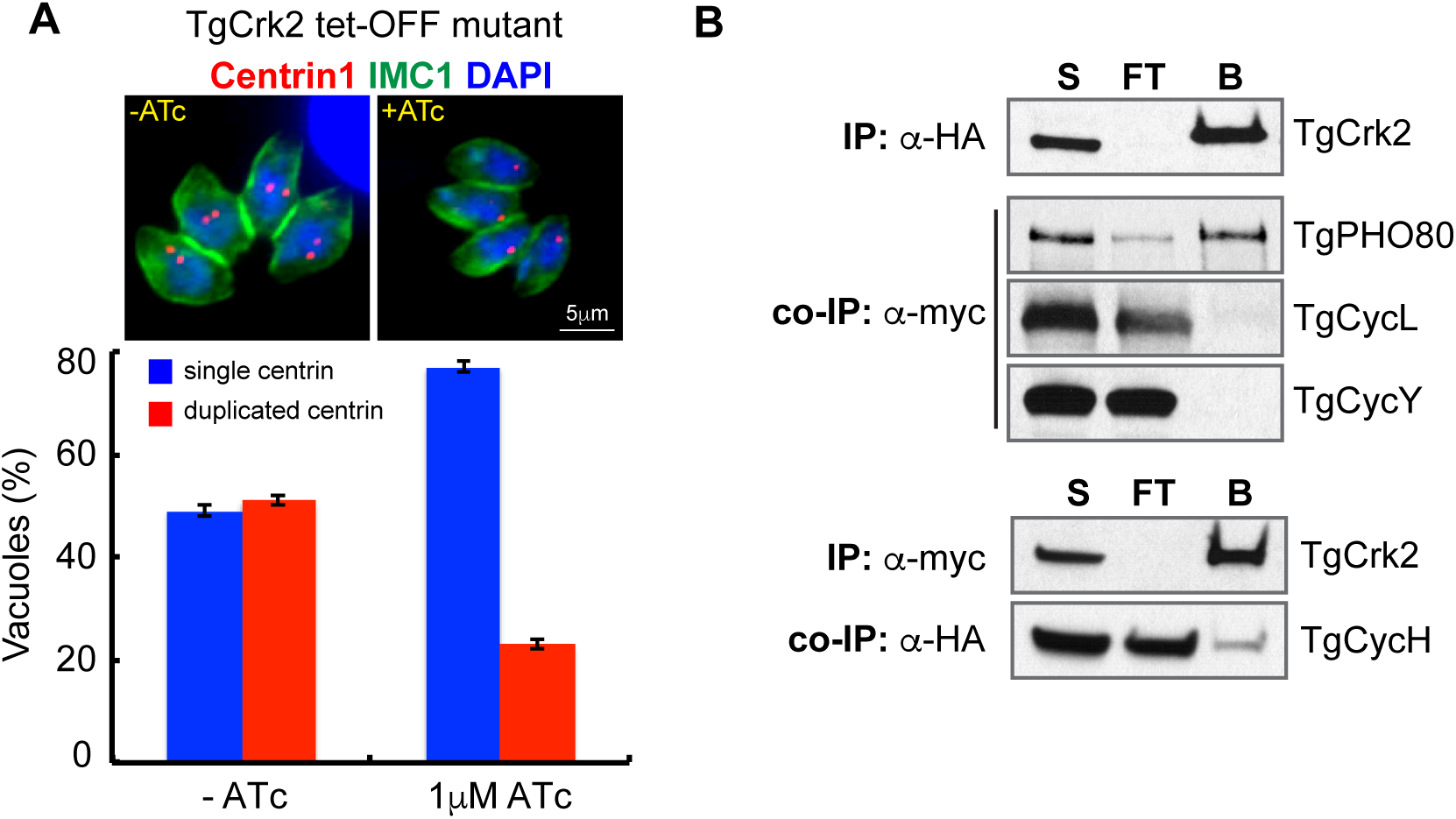
**G1 kinase TgCrk2 predominantly interacts with cytoplasmic TgPHO80 cyclin**. (A) Downregulation of TgCrk2 led to growth arrest in G1 phase prior to centrosome duplication. The TgCrk2 tet-OFF mutant was grown for 24 h with or without 1µg/ml ATc, and co-stained with α-human Centrin1 (red, centrosome), α-IMC1 (green, parasite cytoskeleton and internal buds) and DAPI (blue, nucleus). Vacuoles with duplicated (red column) or single centrosomes (blue column) were randomly selected and counted in -ATc and +ATc populations. The average numbers and standard deviations of three independent experiments are plotted. (B) TgCrk2/cyclin complexes were immunoisolated from the soluble fraction [S] of parasites co-expressing endogenous TgCrk2^HA^ and ectopic myc-tagged TgPHO80, TgCycL, TgCycY (upper panels), and endogenous TgCycH^HA^ co-expressed with ectopic allele of myc-tagged TgCrk2 (lower panels). Ectopic expression was regulated by destabilization domain (DD). Beads with precipitated complexes [B] and depleted soluble fraction [FT] were probed with α-myc and α-HA antibodies to detect cyclins and to confirm efficient pulldown of TgCrk2 (the IP panels on the top). TgCrk2 formed stable complexes with TgPHO80 cyclin, and showed weak interaction with TgCycH, while no complexes were detected with TgCycL or TgCycY, confirming specificity of interactions.

### TgCrk1 and its cyclin partner TgCycL regulate assembly of the daughter scaffold

Conditional loss of TgCrk1 in the tet-OFF mutant resulted in severe chromosome mis-segregation and accumulation of deformed zoites (Fig 2). To more precisely define the TgCrk1-loss phenotype, we focused on the early mitotic steps just prior to daughter bud formation. The MORN1 protein associates with two compartments that are duplicated in mitosis, the spindle compartment (centrocone) and basal complex referred to here as the MORN-ring (Fig 5A, image a) [27, 28]. Utilizing the MORN1 marker, we determined that in parasites lacking TgCrk1 (+ATc for 16 h) the development of the daughter MORN1 rings was defective (Fig 5A, compare image a to c, e). Deficiency of the basal complex became more evident when newly produced alveolar sacs (Fig 5A, image f) accumulated near fragmented MORN-rings in the late stages of mitosis (Fig 5A image e). In addition, the normal coordination of mitosis with cytokinesis failed and the IMC compartment appeared as an unstructured mass (Fig 5A, image h). In addition, TgCrk1 deficiency affected the structural integrity of the tachyzoite apical end (Fig 5B). Following ATc treatment the robust cone-shaped apical cap (ISP1 staining) [47] became a weak rod-like or lopsided structure that caps a deformed IMC1-positive mass (Fig 5B). Despite severe defects in mitosis caused by the loss of TgCrk1, the duplication and segregation of centromeres, centrosomes and the plastid (S4A Fig) and nuclear division were evident (Fig 5A, image j). Altogether these results indicate that TgCrk1 primarily regulates cytokinesis but not karyokinesis.

**Fig 5.**
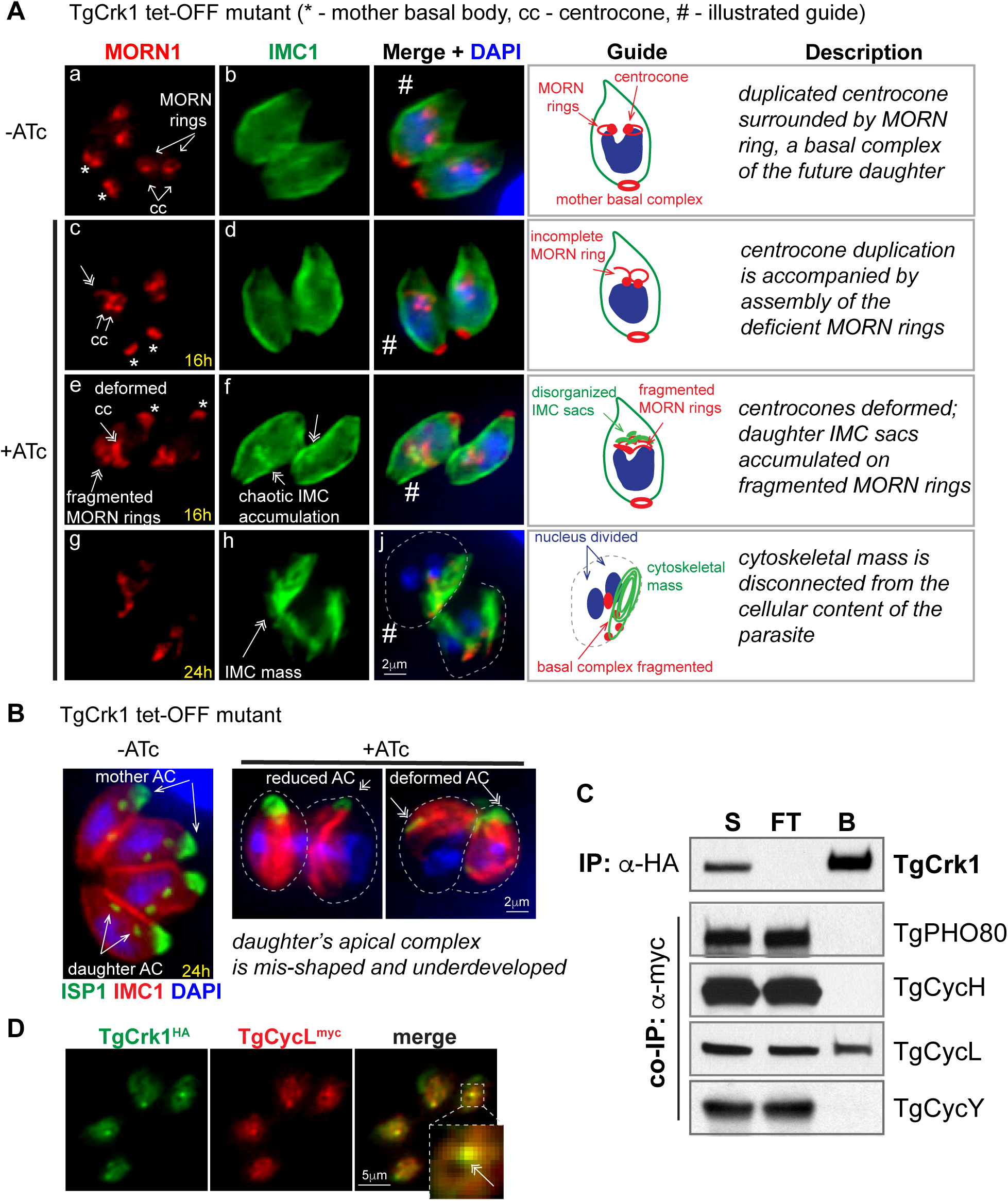
**Loss of TgCrk1 that forms a stable sub-nuclear complex with TgCycL leads to abnormal assembly of the daughter cytoskeleton**. (A) Cytological defects of TgCrk1 tet-OFF mutant were analyzed using antibodies against the basal complex marker MORN1 (red), cytoskeletal marker IMC1 (green) and nuclear staining DAPI (blue). The guide panel shows schematics of the parasites captured in IFA images (#) and parasite features are summarized in the description panel. During normal mitosis depicted in the -ATc images, membrane protein MORN1 localizes to the mother basal complex, intranuclear spindle compartment centrocone, and to the attached MORN rings (image a). Downregulation of TgCrk1 with 1 µg/ml ATc for 16 h resulted in incomplete assembly (image c) or fragmentation (image e) of the MORN rings and associated accumulation of IMC sacs (image f). Fully arrested vacuoles shown in the bottom panel (24 h +ATc) exhibit a catastrophic phenotype; the chaotic daughter cytoskeletal mass (image h) disconnected from parasite cytoplasm. (B) TgCrk1 tet-OFF mutant was stained with α-ISP1 antibodies to monitor changes in the apical complex. Image on the left illustrates proper expression and localization of the marker in S/M tachyzoites (-ATc). Downregulation of TgCrk1 (+ATc) caused severe morphological changes in the apical cone and reduced expression of ISP1 protein. Dotted line indicates boundary of the mis-shaped parasite. Major abnormalities are labeled and indicated by double-headed arrows. (C) TgCrk1-TgCyclin complexes were immunoisolated from parasites co-expressing TgCrk1^HA^ and myc-tagged TgPHO80, TgCycL, TgCycY and TgCycH (used transgenic strains are listed in S1 Table). The soluble fraction before [S] and after immunoprecipitation [FT], and protein complexes on the beads [B] were probed with α-myc antibody to detect TgCyclins and with α-HA antibody to verify TgCrk1 pulldown (top panel). The results revealed a dominant TgCrk1-TgCycL complex. (D) Dual-tagged parasites expressing TgCrk1^HA^ and TgCycL^myc^ were stained with a α-HA (green) and α-myc (red) antibodies. Proteins display similar localization patterns with particular accumulation in the nuclear sub-compartment (insert, arrow).

Constitutively expressed TgCrk1^Ty^ (S4B Fig) and TgCycL^HA^ are similarly localized and knockdown phenotypes of TgCycL and TgCrk1 tet-OFF mutant parasites were also similar (Fig 2 and Fig 3), indicating these two proteins may be paired in *T. gondii*. Indeed, co-IP experiments using the dual epitope-tagging approach employed with TgCrk2 above, confirmed preferential interaction of TgCrk1 with TgCycL (Fig 5C). Furthermore, IFA of the dual-tagged strain co-expressing TgCrk1^HA^ and TgCycL^myc^ revealed tight co-localization of these factors in a unique sub-nuclear compartment (Fig 5D) that was independent of the nucleolus (TgNF3, [48]), centromere-compartment (TgCenH3, [23]), centrocone (MORN1, [26]), or nascent particles containing the RNA polymerase complex (TgRPB4, [49]) (S4C Fig).

### Apicomplexa-specific TgCrk4 and TgCrk6 regulate mitosis in tachyzoites

Conditional knockdown of TgCrk4 and TgCrk6 tet-OFF mutants also caused severe mitotic defects (Fig 2) indicating that similar to TgCrk1, these Apicomplexa-specific kinases function in the second half of the tachyzoite cell cycle. IFA analysis of endogenously tagged TgCrk6^HA^ and TgCrk4^HA^ determined that these proteins have different subcellular localization (Fig 2). TgCrk4^HA^ was distributed in large cytoplasmic aggregates with accumulation in the apical perinuclear region (S2C Fig), while TgCrk6^HA^ extended its nuclear localization to the centromeric region (S2D Fig, CenH3 and Centrin1 staining) [23].

To build clues to TgCrk6 function, we performed detailed IFA analysis following short term ATc treatment (16 h) using the MORN1 (mitotic structures) and IMC1 (cytoskeleton) cell cycle markers. The normal assembly of the daughter scaffold is initiated in late S phase followed by duplication/separation of the centrocone (Fig 6A, -ATc) [2, 3]. Cytokinesis progressed in TgCrk6-deficient parasites, yet the centrocone spindle compartment was not properly duplicated as evidenced by the single MORN1-positive dot positioned between two growing daughter buds (Fig 6A, +ATc). Other evidence supported TgCrk6 function in karyokinetic processes. ATc-downregulation of TgCrk6 disrupted the usual dynamics of kinetochores visualized by co-staining of the kinetochore complex component, TgNdc80^myc^, and acetylated Tubulin A that labels active sites of the microtubule assembly including spindle and internal daughters (Fig 6B) [24]. In normal parasites, the TgNdc80^myc^ signal largely disappeared at mid-bud development (Fig 6B, -ATc image e). By contrast, TgCrk6 deficient parasites (+ATc) retained single assembled kinetochores well into the budding process (Fig 6B, image g). Longer term ATc incubations (>24 h) amplified the loss of coordination between cytokinesis and karyokinesis leading to the catastrophic phenotype of severe DNA mis-segregation and assembly of buds lacking DNA shown in Fig 2. These results combined with the observed accumulation of the nuclear TgCrk6^HA^ in the centromeric region during peak expression in S/M phase (S2D Fig) supported a key role for TgCrk6 in *T. gondii* spindle regulation.

**Fig 6.**
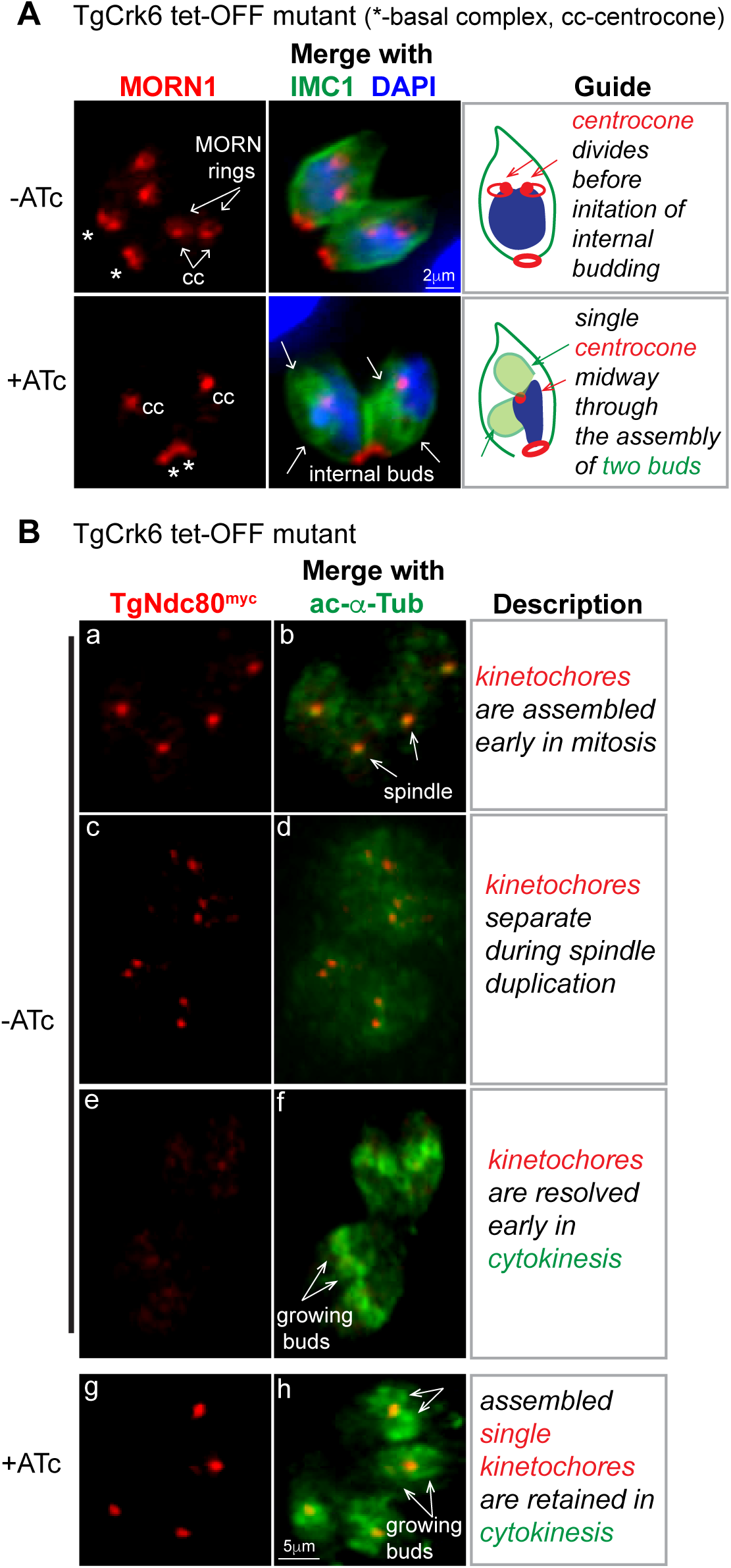
**Nuclear TgCrk6 regulates spindle biology in tachyzoites**. (A) TgCrk6 tet-OFF mutant parasites were grown in the absence (upper row) or presence of 1 µg/ml ATc for 16 h (bottom row) and analyzed by IFA using α-MORN1 (red, centrocone and basal complex) and α-IMC1 (green, parasite shape and internal buds) antibody. Chromosome dynamics was detected by DAPI staining (blue). Downregulation of TgCrk6 (+ATc) led to an inability of the centrocone compartment to duplicate during mitosis. The guide panel includes a cartoon and a description of the analyzed structures and observed deficiencies. (B) The kinetochore dynamics was analyzed in the TgCrk6 tet-OFF mutant parasites co-expressing TgNdc80^myc^ using α-myc antibody. To identify vacuoles in cytokinesis, microtubules of the growing internal buds were stained with antibody against acetylated Tubulin A (green). Kinetochore dynamics in TgCrk6 tet-OFF mutant after 24 h treatment with or without 1µg/ml ATc is summarized in the guide panel on the right. TgCrk6-deficient parasites (+ATc) retained a single assembled kinetochore (image g) positioned between two internal daughters (image h).

Similar to TgCrk6, knockdown of TgCrk4 caused defects in mitosis (Fig 2), although TgCrk4 differed from TgCrk6 in being localized to the cytoplasm rather than the nucleus (Fig S2B versus S2D Fig). This difference led us to examine the role of TgCrk4 in regulating the cytoplasmic components of the mitotic machinery. Asexual stages of *T. gondii* divide by enclosed mitosis (as do most apicomplexans) that coordinates attachment of nuclear centromeres to kinetochores/spindle and to a unique centrosome containing two independent functioning core structures [3, 22]. Consistent with a role in controlling mitosis through cytoplasmic structures, down regulation of TgCrk4 with 1µg/ml ATc led to defective duplication of both centrosomal cores (Centrin1/outer core and CEP250^myc^/inner core), but did not affect centromere duplication/segregation (CenH3 marker) or nuclear division (Fig 7A). Interesting, plastid segregation, which is controlled by the centrosome [50], was also defective in parasites lacking TgCrk4 (Fig 7A, TgAtrx1 marker). Although TgCrk4-deficient parasites showed abnormal centrosome replication (under and over reduplication) (Fig 7A), we did not observe uncoupling of the centrosome cores (Fig 7B) as we have documented in some temperature sensitive mutants [3]. Moreover, we found that centrosome re-duplication occurred around assembled kinetochores (Fig 7C) despite the disruption in normal centrosome stoichiometry. The proper ratios of centrosome to kinetochore observed in a regular tachyzoite mitosis are established (Fig 7C, -ATc; 2:1 images a, b; 2:2 images c, d; 2:0 images e, f). By contrast, down regulation of TgCrk4 led to abnormal stoichiometry of 4 centrosomes to 2 assembled kinetochores (Fig 7C, +ATc, images g, h). Further examination revealed that only one of the reduplicated centrosomes remained associated with the nucleus (Fig 7F, inset), which may explain why the loss of TgCrk4 did not lead to unregulated karyokinesis. Altogether, our results support the role of TgCrk4 kinase in the regulation of centrosome duplication and segregation during mitosis within the context of other essential mitotic regulatory controls such as TgCrk6 above.

**Fig 7.**
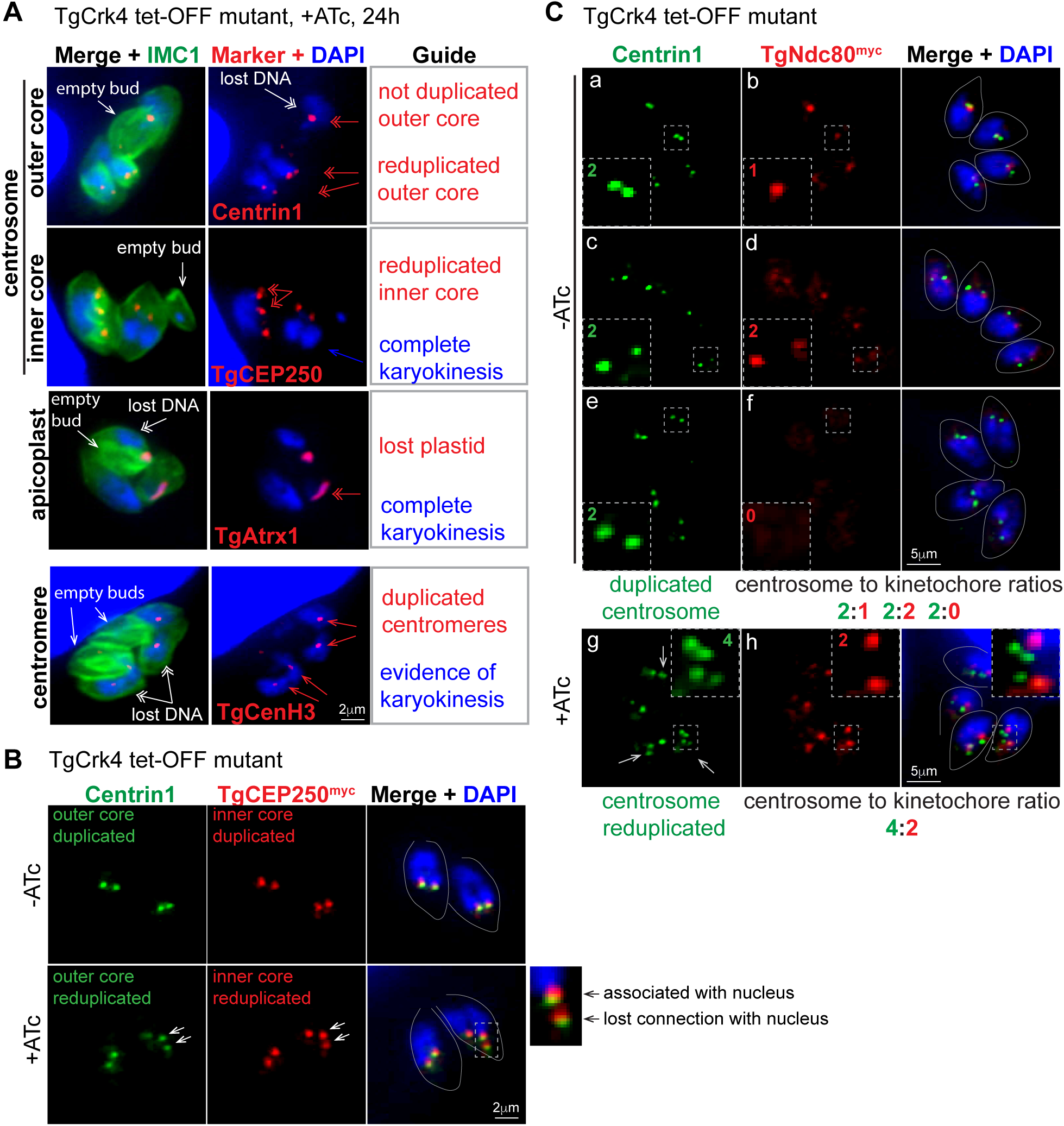
**Loss of cytoplasmic TgCrk4 leads to abnormal centrosome duplication**. (A) TgCrk4 deficiency affected duplication of the structures localized in the cytoplasm. TgCrk4 tet-OFF mutant parasites were treated with 1µg/ml ATc for 24 h and stained with α-IMC1 (surface), DAPI (DNA) and the markers of the following structures: centromere (α-TgCenH3), outer core of the centrosome (α-Centrin1) and apicoplast (α-TgAtrx1). To visualize inner core of centrosome, we introduced recombinant TgCEP250^myc^ protein on the fosmid in TgCrk4 tet-OFF mutant and stained parasites with α-myc antibody. Dynamics of organelles and structures are indicated with red arrows and summarized in the Guide panel. General deficiencies caused by the loss of TgCrk4 are indicated with white arrows. (B) The reduplicated centrosome of TgCrk4-deficient parasites preserved internal integrity. TgCrk4 tet-OFF mutant parasites co-expressing TgCEP250^myc^ were stained with α-Centrin1 (outer centrosomal core) and α-myc (TgCEP250^myc^, inner centrosomal core) after 16 h growth with or without 1µg/ml ATc. White arrows indicate re-duplication of both centrosome cores in TgCrk4 deficient parasite. The enlarged merged image shows that only one of the centrosomes remains connected to the nucleus. (C) Reduplicated centrosomes are associated with assembled kinetochores in TgCrk4 deficient parasites. Transgenic TgCrk4 tet-OFF mutant expressing kinetochore marker TgNdc80^myc^ was grown with or without 1µg/ml ATc for 16 h and co-stained with α-Centrin1 (centrosome), α-myc (TgNdc80^myc^, kinetochore), and DAPI. In mitotic cells expressing TgCrk4 (-ATc), centrosome duplication and kinetochore assembly occurs once per chromosome cycle. Kinetochores are assembled after centrosome segregation (2 centrosomes:1 kinetochore, images a, b), duplicated after the spindle break (2 centrosomes: 2 kinetochores, images c, d) and segregated/disassembled early in the budding (2 centrosomes: 0 kinetochores, images e, f). Lower panel (+ATc) shows vacuole of four TgCrk4-deficient parasites with three cells (white arrows) containing reduplicated centrosomes associated with assembled kinetochores (4 centrosomes: 2 kinetochores, images g, h). Number of structures per parasite is shown in the each insert. Parasite shapes are outlined in the merge panel.

Interestingly, TgCrk4 and TgCrk6 were cyclically expressed during tachyzoite replication (S2 Fig) with the peak expression in S/M phase consistent with functions in regulating mitotic processes. In higher eukaryotes, mitotic Cdk activity is typically controlled by an oscillating cyclin partner, while the Cdk protein is constitutive [29, 38, 39]. To identify the cyclin partners for TgCrk4 and TgCrk6, we performed co-IPs from dual tagged strains expressing TgCrk4^HA^ or TgCrk6^HA^ and four different ectopically expressed TgCyclins (See Material and Methods). No detectable interaction between the TgCyclins tested and TgCrk4 and TgCrk6 was observed (S3D Fig) suggesting *T. gondii* may have become dependent on direct mechanisms of dynamic expression to regulate mitotic TgCrk4 and TgCrk6 factors leading to the loss of a periodic activating cyclin partner.

## DISCUSSION

The molecular basis of cell cycle regulation in eukaryotes has been mainly shaped by studies in one branch of eukaryotes, Unikonta that includes the clades animalia, fungi and amoebas [29]. This is a significant deficiency because the replication biology of eukaryotes from the Bikonta branch, comprised of the three supergroups, the Excavata, SAR (<u>S</u>tramenopiles, <u>A</u>lveolates and <u>R</u>hizaria) and <u>A</u>rchaeplastida, is quite extraordinary, if also beyond our reach experimentally [51–53]. From this point of view, our genetic analysis of *T. gondii* Cdk-related kinases and cyclins provides much needed insight into the cell cycle regulation of an ancient protozoan from the SAR supergroup. One of the core findings of this study is the surprising complexity and unusual regulation of cell cycle controls that are essential for Apicomplexa cell division. The results of multiple gene knockouts in higher eukaryotes reveals that a single active Cdk (Cdk1/2 family) is sufficient to sustain basic chromosome segregation in the somatic cells of both multicellular and unicellular eukaryotes [29, 54]. By contrast, this study and a second project in progress (TgCrk5 studies, Naumov and White, personal communication) have established that *T. gondii* requires five Crks to successfully regulate the peculiar parasite cell cycle called endodyogeny (Fig 8). Two Crks regulate centrosome duplication (TgCrk4) and organization of the daughter bud cytoskeleton (TgCrk1) during interwoven S, M and C phases. *T. gondii* has also evolved independent controls for the restriction or START checkpoint in G1 (TgCrk2), the DNA licensing checkpoint in S phase (TgCrk5, Naumov and White, personal communication) and the spindle assembly checkpoint (TgCrk6) acting at the metaphase to anaphase transition in mitosis (Fig 8). The high number of the putative cell cycle checkpoints was unexpected and this favors a model of much tighter cell cycle regulation in the Apicomplexa than previously thought [1]. In fact, the lack of reversible and abundant catastrophic phenotypes observed in *T. gondii* cell cycle mutants [9], which was often interpreted as a lack of cell cycle controls, is likely a consequence of the complexity of this system. We propose that apicomplexan parasites evolved separate Crks for individual cell cycle stages to facilitate switches between flexible division modes (refs). During the chromosome cycle of schizogony or endopolygeny, the G1 phase is completed only once and is uncoupled from the multiple rounds of the S/M phase, which are, in turn, uncoupled from the budding process until the very last unified S/M/C phase. Therefore, evolution of multiple Crks offers independent control of the segments, permitting modular regulation of the complex cell cycles, yet, leaving an open question of the master regulator(s) of the switch.

**Fig 8.**
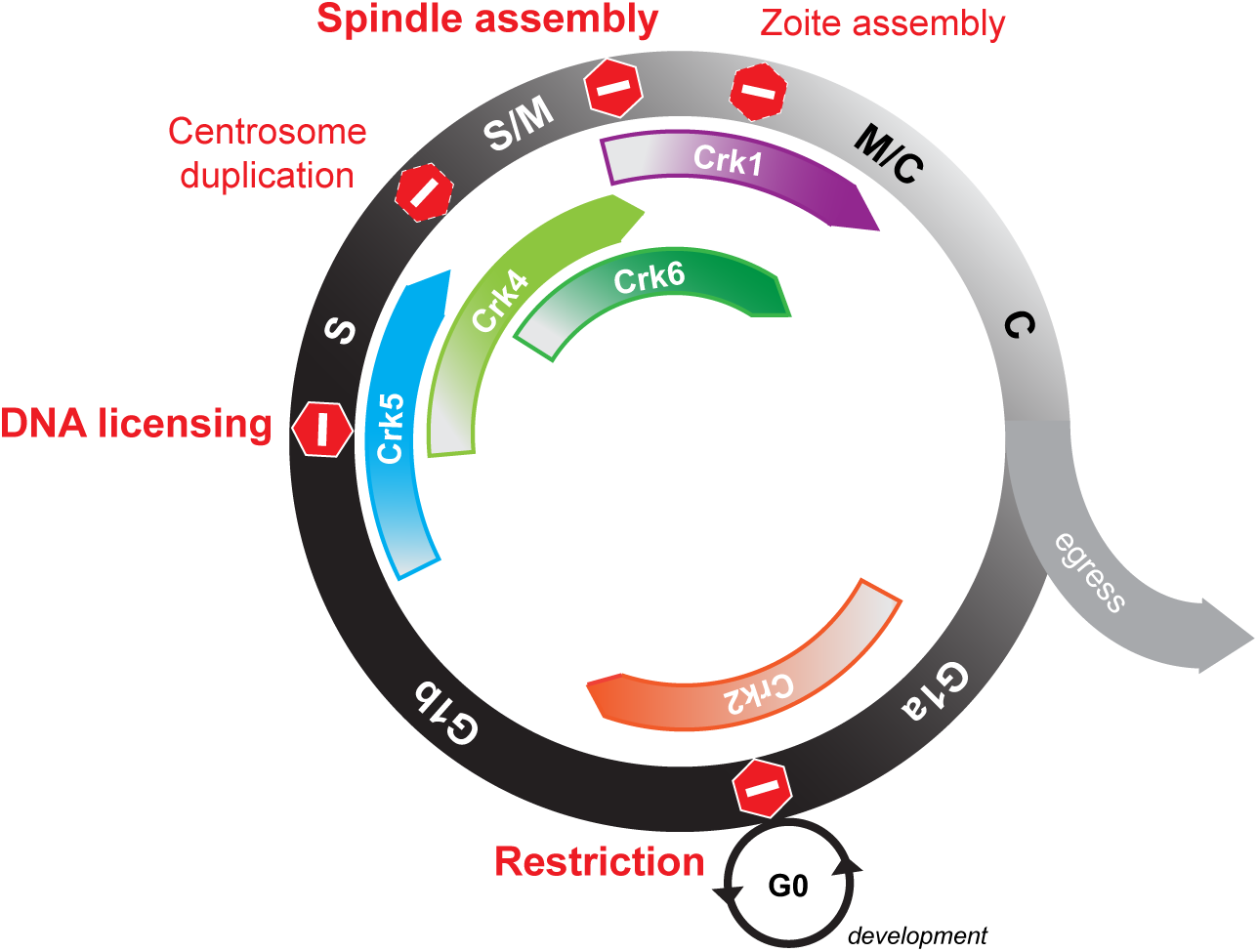
**A roadmap of putative checkpoints of the tachyzoite cell cycle**. Cartoon summaries findings of the current study. The tachyzoite cell cycle consists of G1, S-phase, and mitosis that overlaps and is coordinated with cytokinesis (Black and white circle). Binary division of tachyzoites continues for 5-6 rounds within the same vacuole, and then, parasites lyse the host cell and egress. We identified several potential stopping points (checkpoints) regulated by Cdk-related kinases of *T. gondii*. Red hexagons indicate the timing of the synchronized growth arrest or retardation caused by dis-regulation of the major cellular pathway. It appears that *T. gondii* retained three conserved stop points in the cell cycle. The restriction point in G1 that is related to cell differentiation and dormancy and is regulated by TgCrk2 kinase in complex with TgPHO80 cyclin (orange arrow). Licensing of DNA replication is likely under control of atypical TgCrk5 (blue arrow) (Naumov and White, personal communication), while novel TgCrk6 (dark green arrow) might operate the spindle assembly checkpoint in metaphase. Specialized mitosis of apicomplexan parasites seems to acquire two additional points of cell cycle control. We showed that coccidian-specific kinase TgCrk4 was required to maintain proper stoichiometry of the novel bipartite centrosome (light green arrow). And distantly related to higher eukaryotic Cdk11, TgCrk1 appears to control a vital parasite-specific process of the zoite assembly (purple arrow).

Analyzing the *T. gondii* cell cycle we noticed multiple parallels in topology of the cell cycle regulation between apicomplexans and a few studied Bikonta models, particularly, plants. First, similar to plants, mitosis in *T. gondii* is regulated by clade-specific Cdks (Fig 1) [55]. Second, mitotic TgCrk4, TgCrk5 (Naumov and White, personal communication) and TgCrk6 are dynamically expressed (protein and mRNA), which is a distinctive feature of the CDKB family kinases in plants [55, 56]. Third, similar to most Archaeplastida members, apicomplexan parasites do not posses or encode a highly diverged Cdc25 phosphatase ortholog (EupathDB) [56, 57], which primary function is to activate mitotic Cdks. Interestingly, based on the functional parallels between *Arabidopsis thaliana* CDKB1;1 and *Drosophila melanogaster* Cdc25, plant biologists proposed that the Cdc25-controlled onset of mitosis may have been evolutionary replaced by plant-specific B-type Cdk pathway [57]. Whether the similar scenario had taken place in Apicomplexa evolution will require further studies, involving broader analysis of the Bikonta organisms. Unfortunately, the current limitation of the bikont studies also does not permit us to make a definitive conclusion of whether the plant-like features of the Apicomplexan cell cycle were inherited at the time of the Unikonta and Bikonta diversion or were the result of a secondary symbiosis of the Chromoalveolata and red algae [58].

The majority of characterized cell cycle cyclin-Cdk complexes are composed of constitutive and dynamic subunits with the cyclin generally the oscillating partner [38]. Consistent with the concept that functional topology is more important than the conservation of individual parts [59], we found that *T. gondii* cyclins differed from their high eukaryotic counterparts in being pre-dominantly constitutively expressed. Intriguingly, a single oscillating cycin, TgCycY, was not essential for tachyzoite division nor did it interact with the seven TgCrks we analyzed (S3D Fig) and, therefore, a possible Cdk-independent role will need to be explored in future studies [60]. In fact, only one cell cycle TgCrk2-TgPHO80 complex was detected in which both subunits seem to be constitutively expressed in tachyzoites, suggesting mechanisms other then cyclin-binding that regulate TgCrk2 activity in G1 phase. On the contrary, mitotic TgCrk4 and TgCrk6 are rare examples where Cdk-related kinases are dynamically expressed and not found in the complex with TgCyclins (S3D Fig). Given that the role of cyclin is to provide a temporal context to Cdk function, it is possible that *T. gondii* no longer needs cycling partners for TgCrk4 and TgCrk6. There is reason to speculate that constitutively expressing mitotic kinases might not be ideal given the complexity of the apicomplexan mitosis that is associated with the extensive *de novo* biosynthesis of the motility and invasion apparatus of daughter parasites [4]. Delivering the master conductors "just-in-time" would avoid accidentally triggering the cascade of mechanisms that unfold during mitosis and cytokinesis before the parasite is ready to egress. This hypothesis, however, cannot explain the absence of cyclin interaction with the constitutively expressed TgCrk3 and TgCrk8 (S3D Fig). It is possible that a non-cyclin factor acts as an oscillating component in the complex or, likely, the general metabolic functions regulated by these kinases are needed during all the stages of division. It will require unbiased approaches to identify protein interactions in order to understand how they function.

Another core finding was the strong molecular support for the remarkable physical partitioning of Apicomplexan cell division functions. Similar to other apicomplexans, *T. gondii* divides by enclosed mitosis where a set of tethered structures localized in the nucleus or in the cytoplasm must be constructed/deconstructed (kinetochores, spindle microtubules and striated fibers), duplicated/segregated (bipartite centrosome and centrocone) or assembled (buds) in a timely manner to produce infectious progeny [2, 3, 8]. Moreover, in our previous study [3], we demonstrated that *T. gondii* has divided the regulatory responsibility for karyokinesis and cytokinesis between two unique centrosome cores that have fixed orientation to nuclear and cytoplasmic biosynthetic events. Results of our study here largely support this nuclear/cytoplasmic organization, which was likely needed to overcome limitations of enclosed mitosis. *T. gondii* has evolved the nuclear TgCrk6 mechanism to control events requiring the intranuclear spindle, while cytoplasmic TgCrk4 regulates centrosome duplication and associated plastid segregation. While proper assembly of the nuclear spindle and cytoplasmic daughter bud were essential processes, reduplication of the centrosome in TgCrk4-deficient parasites only partially affected tachyzoite survival. We believe that apicomplexans may have relaxed the control of centrosome reduplication in order to more easily adapt cell division to the scale needed in different hosts [3, 22]. It should be noted that dissolution of the nuclear membrane during open mitosis in higher eukaryotes is a key difference with apicomplexan cell division that may permit extensive re-arrangements of the nuclear (e. g. kinetochore, spindle) and cytoplasmic (e. g. centrosome) structures by a single Cdk.

A possible exception to the nuclear/cytoplasmic functional organization is the assembly of the cytoplasmic daughter buds that was unexpectedly controlled by the nuclear TgCrk1-TgCycL complex. Initiation of the daughter buds near the centrocone, spindle pole, which continues to grow with progression of mitosis, is a specialized event occurring in the budding cycle of Apicomplexa [2, 25]. How does the nuclear TgCrk1 regulate assembly of a cytoplasmic structure? Recent studies of eukaryotic splicing kinase Cdk11 discovered an unexpected role for this kinase in mitotic progression [61]. It has been shown that activity of Cdk11 is required to regulate sister chromatid cohesion [62]. Since knockdown of *T. gondii* TgCrk1 did not affect DNA segregation, it is possible that the role of TgCrk1 was re-adapted to regulate splicing of mRNAs whose products will be required to control assembly of the daughter buds. Our hypothesis is supported by the fact that many components of the cytoskeleton are delivered “just-in-time” during cell cycle progression [4]. In future studies, the analysis of transcriptome changes caused by TgCrk1 deficiency will help determine whether this kinase operates primarily as a regulator of mRNA expression.

Transmission stages are formed at the end of each apicomplexan life cycle, that are, in most species, specialized G1/G0 states. For example, mature bradyzoites and sporozoites of *T. gondii* remain growth arrested until appropriate external signals from the same or new host trigger recrudescence or de-differentiation, respectively, resulting in re-entry into G1 phase of the asexual proliferative cycle (Fig 8). In higher eukaryotes, Cdk4/6-Cyclin D complexes are responsible for the cell fate decision to divide or differentiate (restriction checkpoint) [63, 64], which are factors not present in the Apicomplexa. Here we showed that *T. gondii* parasites have replaced the canonical G1 machinery of higher eukaryotes with a novel complex of TgCrk2 (Cdk1/2 family) and TgPHO80 cyclin to regulate progression through the tachyzoite G1 phase. Recent studies in kinetoplastids and another apicomplexan, *Plasmodium berghei*, determined that a related G1 Crk and a P-type cyclin regulate developmental stages in these protozoans, which is similar to our discoveries of TgCrk2 function in *T. gondii* (Fig 8) [35, 65, 66]. It is also worth noting that *T. gondii* possesses paralogs of TgCrk2 and TgPHO80 cyclin that are developmentally regulated (ToxoDB, S1 Fig, S3 Fig) opening the possibility that the TgCrk2-TgPHO80 pathway has diverse functions in development and could also be involved in the regulation of drug-induced dormancy.

In conclusion, the systematic approach we have used to analyze the cell cycle machinery in *T.gondii* has opened the way into learning how cell division is regulated in apicomplexan parasites. Our study has also evoked important issues that still need to be addressed. For example, greater numbers of TgCrks requires greater coordination; so is there a master regulator after all? Can the complexity of mitosis coupled to cytokinesis in *T. gondii* division explain a rise of multiple mitotic Crks? What are the mechanisms responsible for periodic Crks, and how are these mechanisms controlled? Does the lessons learned here translate to the exotic mitotic mechanisms in related alveolates [51, 52]? Clearly, there is still much to be done to understand the molecular basis of apicomplexan cell division, fortunately, we now have important new genetic tools to go forward.

## Materials and Methods

### Cell culture and parasite growth analysis

*T. gondii* strains RHΔ*hxgprt [67]*, Tati-RHΔ*ku80* [41], and RHΔ*ku80*Δ*hxgprt* [68] were cultured in human foreskin fibroblasts (HFF) according to published protocols [69]. Viability of transgenic strains was measured in plaque assays as previously described [15]. Monolayers of HFF cells were infected with 150-200 parasites per 35mm dish and individual plaques formed after 6 days were stained with crystal violate and counted. To determine division rates, parasites were counted by IFA using α-IMC1 (surface) antibody and DAPI (nucleus) in 50 randomly selected vacuoles in three biological replicates after 24 hours growth. Statistical significance was calculated using an unpaired T-test and Bonferroni correction (Prism6).

### Generation of transgenic tachyzoite strains

Transgenic strains and primers created in the study are listed in S1 Table.

<u>Endogenous C-terminal tagging</u>. TgCrk and TgCyclin genes were tagged by creating a fusion protein with triple HA-, myc- or Ty- epitopes as previously described [68]. Genomic fragments encompassing the 3'-end of the gene of interest were PCR amplified (primers used are listed in S1 Table) and incorporated into plasmids pLIC-HA_3X_-HXGPRT, pLIC-myc_3X_-DHFR-TS or pLIC-Ty_3X_-HXGPRT via ligation independent cloning (InFusion, Clontech). Linearized constructs were then electroporated into the RHΔ*ku80* strain at a ratio of 5µg DNA per 10^7^ parasites. To establish double-tagged transgenic lines we performed sequential electroporations and selections with alternative drugs. Successful tagging was confirmed by PCR, IFA and Western Blot analysis.

<u>Endogenous tagging using CRISPR/Cas9 technology</u>. To introduce 3xHA-epitope to the C-terminus of TgCycY we modified sgUPRT-CRISPR/Cas9 plasmid [70] and replaced the sgUPRT-RNA with the sgTgCycY-RNA sequence located 60 nt downstream of the STOP codon using Q5 DNA polymerase mutagenesis (New England Biolabs) and primers specific for TgCycY gsRNA (S1 Table). We designed a tagging cassette that included 50 nt prior to the STOP codon of the *TgCycY* gene fused to 3xHA-epitope, chloramphenicol resistance marker and 50 nucleotides of the TgCycY 3’UTR located 70 nt downstream of the predicted CAS9 digestion site (included in sgTgCycY-RNA) (primers used are listed in S1 Table). The amplified cassette and sgTgCycY-CRISPR/Cas9 plasmid were mixed in a 1:1 molar ratio and electoporated into RHΔ*ku80* parasites. Transgenic parasites were selected with 20µM chloramphenicol and analyzed by PCR, IFA and Western Blot analysis.

<u>Endogenous N-terminal tagging and promoter replacement</u>. To build tet-OFF mutants of TgCrks and TgCyclins, the 5’ end of the corresponding gene was amplified (primers used are listed in S1 Table), digested with BglII/XhoI or BamHI/XhoI and ligated into the promoter replacement vectors, ptetO7sag4-myc-DHFR-TS or ptetO7sag4-HA_3X_-DHFR-TS [24, 71, 72]. The vector ptetO7sag4-HA_3X__DHFR-TS was created by epitope swap on the ptetO7sag4-myc_DHFR-TS plasmid (Q5 mutagenesis, New England Biolabs). The resulting TgCrk and TgCyclin tet-OFF constructs were linearized, introduced into the Tati-RHΔ*ku80* [41] strain and selected for pyromethamine resistance. Successful recombination into the locus was verified by PCR and individual transgenic clones were screened by IFA. To test conditional regulation, the tet-OFF mutant clones were grown with or without 1µg/ml ATc and protein expression level of the tet-OFF factors was analyzed by Western Blot analysis and IFA. Double-tagged transgenic lines were established by sequential electroporation and corresponding drug selection.

<u>Ectopic N-terminal tagging and conditional expression</u>. Conditional N-terminal tagging of TgPHO80, TgCycL, TgCycY, TgCrk2 and TgRBP4 was performed as follows. Full-length amplicons flanked with indicated restriction sites were obtained from RHΔ*hxgprt* cDNA libraries (primers used are listed in S1 Table), digested and ligated into the conditional expression vector ptub-DD-myc_3X_-CAT [45, 73]. Resulting constructs were introduced into parasite and selected with 20µM chloramphenicol. Conditional expression of the tagged proteins in transgenic parasites grown with and without 100nM Sheild1 was verified by IFA.

<u>Epitope tagging by fosmid recombination</u>. To create recombination cassettes with alternative epitope tags we replaced a 3xHA epitope encoded in the plasmid pH3CG with a 3xmyc epitope (Q5 mutagenesis, New England Biolabs). The resulting pM3CG plasmid was used to amplify gene-specific tagging cassettes that also included a chloramphenicol resistance marker (*T. gondii* selection) and a gentamycin resistance marker (*E. coli* selection). To epitope tag TgNdc80 (encompassed by the fosmid RHfos08I02), TgCEP250 (encompassed by the fosmid RHfos14B04) and TgCycH (encompassed by the fosmid RHfos20N15) tagging cassettes were introduced into a corresponding fosmids as previously described [3, 74]. Recombined fosmids 08I02-TgNdc80_myc_-CAT, 14B04-TgCEP250_myc_-CAT and 20N15-TgCycH_myc_-CAT were transfected into parasites and cultured under 20µM chloramphenicol selection. Expression of tagged protein was confirmed by IFA with α-myc antibody.

### Immunofluorescence analysis

Monolayers of HFF cells were grown on coverslips and infected with parasites under indicated conditions. Cells were fixed in 4% PFA, permeabilized with 0.25% Triton X-100, blocked in 1%BSA and incubated sequentially with primary and secondary antibody [16]. The following primary antibodies were used: mouse monoclonal α-ISP1 (clone 7E8) [47] and α−Atrx1 (clone 11G8, kindly provided by Dr. Peter Bradley, UCLA) [75], α-TgCenH3 [23] (kindly provided by Dr. Boris Striepen, University of Georgia, Athens), α-acetylated alpha Tubulin (Abcam), rat monoclonal α-HA (clone 3F10, Roche Applied Sciences), rabbit polyclonal α-myc (Cell Signaling Technology), α-Human Centrin 2 [17], α-MORN1 (kindly provided by Dr. Marc-Jan Gubbels, Boston College) [27] and α-IMC1 (kindly provided by Dr. Gary Ward, University of Vermont). Alexa-conjugated secondary antibodies of the different emission wavelengths (Molecular Probes, Thermo Fisher Scientific) were used at a dilution of 1:1000. Stained parasites on the coverslips were mounted with Aqua-mount (Lerner Laboratories), dried overnight at 4°C, and viewed on a Zeiss Axiovert Microscope equipped with 100x objective. Images were collected and processed using Zeiss Zen software and were further processed in Adobe Photoshop CC using linear adjustment when needed.

### Immunoprecipitation assay

Transgenic parasites co-expressing epitope-tagged TgCrks and TgCyclins were grown for 30-32 hours at 37°C. When expression of the factor was conditional, 100nM Shield1 was added for the last 3 hours of incubation. Parasites (3x10^8^) were collected, washed in PBS and lysed in 1xPBS with 0.5% NP-40, 400mM NaCl, protease and phosphatase inhibitors (Thermo Fisher Scientific) on ice for 30 min. Total protein extract obtained by centrifugation at 21,000x*g*, 10 min, 4°C was divided and incubated with α-HA or α-myc magnetic beads (MBL International) for 1 h at room temperature. Isolated protein complexes on the beads were washed three times with the lysis buffer and eluted by heating in the Leamlli sample buffer at 65°C, 10 min. Protein extract before and after pull-down, and purified protein complexes were analyzed by Western blotting.

### Western blot analysis

To prepare samples of the total extracts, parasites were purified by filtering through 3µm polycarbonate filters (EMD Millipore), washed in PBS, re-suspended with Leammli loading dye and lysed at 65°C for 10 min. To analyze individual fractions after immunoprecipitation, an aliquot of the fraction was mixed with Leammli loading dye and heated for 10 min at 65°C. After separation on SDS-PAGE gels, proteins were transferred onto a nitrocellulose membrane and probed with monoclonal α-HA (3F10, Roche Applied Sciences), α-myc (Cell Signaling Technology) and α-Tubulin A (12G10, kindly provided by Dr. Jacek Gaertig, University of Georgia) antibodies. After incubation with secondary HRP-conjugated α-mouse or α-rat antibodies, proteins were visualized by enhanced chemiluminescence detection (PerkinElmer).

### Phylogenetic Analysis

Phylogenetic analysis was performed using Phylogeny.fr (LIRMM, France) using the advanced mode [76, 77]. Because of the lack of cell cycle gene annotation in apicomplexans, we searched the *T. gondii* genome for protein kinases with cyclin binding C-helix and proteins containing a cyclin box showing similarity to mammalian Cdks and cyclins, respectively. Then, to reduce complexity of the analysis we first identified Cdk classes preserved in apicomplexans by comparing Cdk-related kinases of *T. gondii* and Cdks of human cells. Then, the full complement of Crks from *T. gondii*, *P. falciparum*, *Chromera velia* and pre-selected Cdks from *H. sapiens* were compared by first aligning sequences using the MUSCLE algorithm set to 16 iterations. Alignments were further refined with Gblocks0.91b to eliminate poorly aligned regions, although allowing gaps and less strict flanking positions in the final blocks. Relationships between sequences were analyzed using PhyML3.0 set up for 100 bootstraps after gap removal. The final tree was constructed with TreeDyn198.3.

## ACKNOWLEDGMENTS

The authors are grateful to Dr. Michael White (USF) for critical reading, discussions and helpful comments on the manuscript. We also thank Dr. Naumov and Dr. White (USF) for permitting to share their finding on TgCrk5 prior to publication.

